# Molecular Determinants of G Protein Subtype-Specific Coupling to the Angiotensin II Type 1 Receptor

**DOI:** 10.64898/2026.09.30.755740

**Authors:** Akua A. Acheampong, Matthew K. Collins, Amanda R. Keller, Andrew P. Feld, Brandi M. Small, Michael Ritt, Carl-Mikael Suomivuori, Sivaraj Sivaramakrishnan, Laura M. Wingler

**Affiliations:** Department of Pharmacology and Cancer Biology, Duke University, Durham, North Carolina, United States; Department of Genetics, Cell Biology and Development, University of Minnesota, Minneapolis, Minnesota, United States; Department of Computer Science, Institute for Computational and Mathematical Engineering, Stanford University, Stanford, California, United States; Department of Molecular and Cellular Physiology, Department of Structural Biology, Stanford University School of Medicine, Stanford, California, United States

**Keywords:** G protein-coupled receptor, angiotensin, allostery, heterotrimeric G protein, ligand

## Abstract

Many G protein-coupled receptors (GPCRs) are capable of activating multiple heterotrimeric G protein subtypes. The angiotensin II type 1 receptor (AT1R) couples to both Gq and Gi family members, and the two families mediate distinct aspects of the effects of angiotensin II (AngII) on blood pressure and fluid balance. Previous work has pointed to fundamental differences in the strength and ligand-dependence of Gq versus Gi activation by AT1R. Here we resolve key aspects of the mechanisms underlying these subtype-specific patterns of G protein coupling to AT1R. Using cellular biosensors, we show that AT1R activates Gq more efficaciously than Gi. Leveraging mini-G proteins, engineered versions of Gα subunits that diverge only in the C-terminal α5 helix, we identify four residues dispersed throughout the α5 helix that are sufficient to account for these differences. We employ radioligand binding to demonstrate that subtype-specific differences originate at the level of allosteric coupling between ligand and transducer binding, and we use Systematic Protein Affinity Strength Modulation biosensors to show that Gq-specific residues in the α5 helix increase affinity for AngII-activated AT1R. We provide evidence that the α5 helix residue with the greatest effect on coupling makes interactions with AT1R intracellular loop 3 that were not resolved in a previously reported AT1R-Gq cryo-electron microscopy structure. These findings enhance our understanding of how AT1R differentially activates Gq and Gi and may ultimately guide pharmacological approaches to modulate certain G proteins downstream of AT1R and other GPCRs.

**SIGNIFICANCE STATEMENT:** This study identifies four residues within the Gα C-terminal α5 helix that govern subtype-specific patterns of Gq and Gi coupling to the angiotensin II type 1 receptor. Our findings demonstrate that allosteric coupling dictates the differential activation patterns of these two G heterotrimeric proteins, advancing our understanding of transducer selectivity and guiding the development of targeted subtype-selective therapeutics.

## INTRODUCTION

Upon agonist stimulation, a single G protein-coupled receptor (GPCR) can initiate an array of cellular responses by coupling to various intracellular transducer proteins, including heterotrimeric G proteins (Gs, Gi/o, Gq/11, and G12/13), GPCR kinases, and β-arrestins. Each G protein subtype initiates distinct signaling events. For example, following dissociation of the Gα and Gβγ subunits, Gαq activates phospholipase C to trigger downstream calcium signaling, while Gαi inhibits adenylyl cyclase to decrease cyclic AMP production. Of the GPCRs whose G protein selectivity has been systematically profiled, a plurality can couple to multiple G protein subtypes, with Gq and Gi being the most common co-coupling.^1^ However, GPCRs typically display pronounced differences in the relative strength of activation of these two subtypes.^1^

The angiotensin II type 1 receptor (AT1R) exemplifies this pattern of primary and secondary coupling to Gq and Gi, respectively, following stimulation with its endogenous ligand, the octapeptide angiotensin II (AngII). In directly comparable measures of G protein activation, such as Gα and Gβγ dissociation, AT1R activates Gq more efficaciously than Gi.^2^ Many canonical functions of AT1R, including vasoconstriction, result from Gq activation. However, AT1R-Gi signaling mediates important physiological functions of AngII in specific cell types. These include regulation of angiotensinogen production in the liver,^3^ sodium reabsorption in the renal proximal tubules,^4^ calcium channels in the adrenal cortex that in turn regulate aldosterone production,^5,6^ and resting metabolic rate in the arcuate nucleus of the hypothalamus.^7^ AngII activates ERK signaling through Gq in cardiomyocytes but through Gi in cardiac fibroblasts.^8^ Gi coupling is also required for mechanoactivation of β-arrestin through AT1R in the heart.^9^ β-Arrestin signaling, whether activated directly by AT1R following agonist stimulation or in a Gi-dependent manner through AT1R mechanoactivation, has cardioprotective effects.^10–13^

The C-terminal α5 helix of the Gα subunit is a key determinant of the receptor specificity of G proteins. Following ligand-induced allosteric changes in GPCRs, the α5 helix inserts into the intracellular cavity formed by the 7-transmembrane domain of GPCRs. Seminal work demonstrated that swapping the distal four residues of Gαq and Gαi2 was sufficient to alter the specificity of these proteins for many GPCRs.^14^ However, regions outside of the C-terminal α5 helix contribute to the differences in the strength of other GPCR-G protein pairings.^2,15^

Together these data indicate that the determinants of GPCRs’ pattern of G protein coupling are receptor-specific. In this work, we define the mechanisms underlying subtype-specific differences in G protein coupling to AT1R, leveraging the structural information and pharmacological tools available for this receptor. We show that four discrete residues in the Gα α5 helix are sufficient to account for differences in AT1R coupling to Gq versus Gi. These residues are dispersed throughout the last 11 residues of the helix, rather than localized in the distal tip. We further demonstrate that these subtype-specific differences can be traced to the earliest stages of G protein activation, ligand-induced allosteric effects on G protein interactions with AT1R.

## MATERIALS AND METHODS

### Cell culture and transfection

HEK293 suspension cells (Inducible Expi293 Cells, Thermo Fisher Scientific, Waltham, MA, RRID: SCR_008452) were maintained at 37°C in 8% CO_2_ with shaking at 125 rpm. Cells were transiently transfected with plasmids using ExpiFectamine 293 (Thermo Fisher Scientific) or FectoPRO (Sartorius, Göttingen, Germany, RRID: SCR_003935). For ExpiFectamine 293 transfections, cells were transfected at a concentration of 3.0e6 cells/ml with 1 μg DNA/ml of culture. ExpiFectamine 293 Transfection Enhancers were added approximately 18 h post-transfection. For FectoPRO transfections, cells were transfected at a concentration of 3.0e6 cells/ml with 0.75 μg DNA/ml of culture. Sodium valproate (3 mM final concentration) and glucose (0.4% final concentration) were added approximately 18 h post-transfection.

### Radioligand binding

AT1R-miniG constructs for radioligand binding were cloned by genetically fusing human codon-optimized miniGq (miniGs/q70^16^) or miniGi (miniGs/i43^16^) to the C-terminus of AT1R in pcDNA-Zeo-tetO-FLAG-AT1R^17^ (“unfused AT1R,” RRID: Addgene_222856) via a (GGGSG)_2_ linker. Additional mutations in AT1R and mini-G proteins were introduced with the QuikChange Lightning Site-Directed Mutagenesis Kit (Agilent, Santa Clara, CA, RRID: SCR_013575). Cells were transfected with these plasmids as described above. AT1R expression was induced 2 days post-transfection by addition of doxycycline (4 μg/ml) and sodium butyrate (5 mM). At 72 h post transfection, cells were harvested at 500 *g* for 5 min at room temperature, washed once with phosphate-buffered saline (PBS), harvested again, and frozen using liquid nitrogen. Pellets were thawed on ice and resuspended in homogenization buffer (75 mM Tris pH 7.4, 2 mM EDTA, 5 μM leupeptin, 1 mM benzamidine); the suspension was homogenized on ice using a Dounce homogenizer (100 times) and centrifuged at 500 *g* for 5 min at 4°C. The supernatant was transferred to a new tube and centrifuged at 16,100 *g* for 1 h at 4°C. Pellets (crude membrane homogenates) were resuspended in cold resuspension buffer (75 mM Tris pH 7.4, 2 mM EDTA, 5 μM leupeptin, 1 mM benzamidine, 12.5 mM MgCl_2_), aliquoted, and flash frozen in liquid nitrogen for storage at -80°C.

Competition radioligand binding assays were performed to assess the allosteric effects of mini-G proteins on ligand affinity for the receptor. Membranes were incubated in a 96-well, 2-ml deep-well plate (Thermo Fisher Scientific) for 1.5 h at 25°C in assay buffer (50 mM Tris, 150 mM NaCl, 12.5 mM MgCl_2_, 0.2% bovine serum albumin, 5 μM leupeptin, 1 mM benzamidine) with 2.5 nM [^3^H]-olmesartan (10 Ci/mmol, American Radiolabeled Chemicals, St. Louis, MO) and a serial dilution of unlabeled competitor ligand, with a single technical replicate for each condition. Nonspecific binding was determined in the presence of 10 µM unlabeled candesartan, and total binding was determined in the absence of a competitor. Reactions were terminated by rapid vacuum filtration through Whatman GF/B Filter Paper using a Brandel 96-well harvester (Gaithersburg, MD). Filters were washed three times with ice-cold 50 mM Tris pH 7.4 to remove unbound radioligand. Bound radioactivity retained on the filters was quantified using a Packard Tri-Carb 2100TR liquid scintillation analyzer (Meriden, CT).

For saturation binding experiments, membranes were incubated with a titration of [^3^H]-olmesartan in duplicate. The concentration of each [^3^H]-olmesartan dilution was determined empirically by scintillation counting. Candesartan (10 µM) was used to determine non-specific binding at each radioligand concentration.

### SPASM sensors

The SPASM biosensors contain FLAG-human AT1R fused at its C-terminus to the Large BiT (LgBiT) fragment of NanoLuciferase, an ER/K linker, tagRFP, the Small BiT (SmBiT) complementation fragment, and a terminal mini-G protein (miniGq or miniGi) in the pcDNA5/FRT/TO plasmid backbone.^18,19^ The control construct contained all elements except the mini-G protein. HEK293 suspension cells expressing SPASM sensors were harvested 2 days post-transfection (500 *g*, 3 min), and the pellets were washed once in PBS at 100% of original culture volume. The samples were resuspended in GPMV buffer (200 mM NaCl, 0.5 mM MgCl_2_, 5 mM KCl, 0.75 mM CaCl_2_, 100 mM bicine pH 8.5) at 60-70% of original culture volume and were incubated for 2 h at 30°C with shaking at 140 rpm. The samples then underwent three centrifugation steps (1000 *g,* 2 min at 4°C) to pellet and discard cells and debris. The final supernatant was centrifuged (3220 *g*, 40 min at 4°C) and resuspended twice to pellet the GPMVs containing SPASM sensors and remove residual bicine. GPMVs were resuspended in FRET buffer (150 mM NaCl, 5 mM MgCl_2_, 10 mM KCl, 20 mM HEPES pH 7.4) containing 10% glycerol and flash-frozen in liquid nitrogen for storage at -80°C.

All experiments were carried out in FRET buffer. To ensure uniform construct concentration across biological replicates, RFP signals were normalized for each newly prepared batch of GPMVs as follows: stock constructs were diluted 1:5 in FRET buffer and loaded into a black, clear-bottom, 96-well microplate. RFP fluorescence intensities were measured (excitation: 521 nm; emission: 585 nm) alongside assay buffer blanks. To account for background fluorescence, the mean signal of the assay buffer was subtracted from the mean signal of each individual construct, and the construct exhibiting the lowest RFP signal was used as the reference. In subsequent experiments, the concentrations of all GPMVs in the batch were adjusted to match the fluorescence of the construct with the lowest specific fluorescence. GPMVs were incubated with 2.78 µM coelenterazine-H (Research Products International, Mt. Prospect, IL) for 5 min, and luminescence was recorded (CLARIOstar, BMG Labtech, Cary, NC) for 5 min at 1-min intervals. AngII (10 µM) was added to each well (3 technical replicates per condition), and luminescence was monitored for an additional 30 min.

### G protein dissociation assays

Plasmids for the TRUPATH system were a gift from Bryan Roth (Addgene Kit #1000000163). For the G protein activation assays, cells were co-transfected with a 4:1:1:1 ratio of pcDNA-Zeo-tetO-Flag-AT1R, Gα-RLuc8 (mutant *Renilla* luciferase) (Gαq-RLuc8, RRID: Addgene_140982; Gαi2-RLuc8, RRID: Addgene_140974), Gβ3 (RRID: Addgene_140988), and Gγ9-GFP2 (mutant green fluorescent protein) (RRID: Addgene_140991). AT1R expression was induced 2 days post-transfection by addition of doxycycline (4 μg/ml) and sodium butyrate (5 mM). Cells were harvested 3 days post-transfection and resuspended in assay buffer (Hanks’ balanced salt solution with 20 mM HEPES pH 7.4). Cells were then dispensed into opaque white 96-well plates and incubated with 1.3 µM Prolume Purple (methoxy e-Coelenterazine) (Nanolight Technologies, Prolume, LTD., Pinetop, AZ) for 5 minutes, after which a serial dilution of AT1R ligand in assay buffer containing 0.1% bovine serum albumin was added. Three technical replicates were included for each condition. Plates were read for 6 cycles on the CLARIOstar (BMG Labtech) set at 37°C with 410 nm and 515 nm emission filters. Data from the fourth read (10 min post-ligand addition) were used to calculate the bioluminescence resonance energy transfer (BRET) ratio using the ratio of the emission at 510 nm (GFP2) to the emission at 400 nm (RLuc8).

### Mini-G protein recruitment assays

Wild-type AT1R was genetically fused to RLuc8 via a TGGGGS linker (AT1R-RLuc8) and utilized for all mini-G protein recruitment assays. MiniGs/q71-mVenus and miniGs/i43-mVenus constructs were previously reported.^20^ MiniGs/q71, which contains additional Gαq-specific residues outside of the α5 helix,^16^ was replaced with miniGs/q70 for this study. Cells were co-transfected with a 1:3 ratio of AT1R-RLuc8 to miniG-mVenus. The remainder of the protocol was identical to the G protein dissociation assays described above, except luminescence was monitored at 480 nm (RLuc8) and 530 nm (mVenus).

### Data analysis

The experiments reported in this manuscript were exploratory in nature and not designed to test a prespecified null hypothesis. In all experiments, non-overlapping 95% confidence intervals (CIs) were considered to be statistically significant.

For competition radioligand binding assays, a two-site competition binding model was fit to the data for AT1R-miniG constructs in GraphPad Prism v. 11.0.2 (RRID:SCR_002798), while the optimal model for AT1R unfused was determined by a comparison of fits between one-site and two-site competition models. After normalizing individual curves from each experiment to their internal top and bottom values (0-100%), Area Under the Curve (AUC) values for each curve were determined in GraphPad Prism. The AUC values from independent biological replicates were pooled and then normalized to AT1R unfused (0%) and AT1R-miniGq (100%) to determine the relative Δ integral. Reported relative Δ integral values represent the mean and 95% CIs determined from ≥3 independent experiments.

For saturation radioligand binding experiments, a one-site saturation binding model was fit to the data from each individual experiment in GraphPad Prism using the experimentally derived concentrations of [^3^H]-olmesartan. Reported dissociation constant (Kd) values represent the mean and 95% CIs determined from three independent experiments.

For SPASM sensor experiments, raw luminescence data from triplicate wells were averaged per construct across 5 pre-read cycles to generate a 3-point moving average. After subtracting the mean background signal, each post-ligand addition replicate value (cycles 6-35) was normalized to the final averaged pre-read value for that construct. The AUC was determined for each kinetic course in GraphPad Prism, and the -AngII AUC was subtracted from the +AngII AUC to yield a Δ integral value for each construct. Δ Integral values were normalized to miniGq (100%) to obtain the relative Δ integral. Reported relative Δ integral values represent the mean and 95% CIs determined from ≥3 independent experiments.

For both G protein dissociation and mini-G protein recruitment assays, three-parameter nonlinear regression curves [log(agonist) vs response] were fit to BRET data. BRET data from each experiment were normalized to the top and bottom of the Gαq or miniGq dose-response curve. Fits of these normalized data from independent experiments were used to determine maximal efficacy (E_max_) and potency (logEC_50_) values for each construct. The reported E_max_ and logEC_50_ represent the mean and 95% CIs determined from ≥3 independent experiments.

## RESULTS

### The C-terminal α5 helix of Gα encodes subtype-specific coupling to AT1R

To compare the ability of AT1R to activate Gq and Gi, we used a BRET-based assay that monitors dissociation of the Gα and Gβγ subunits.^21,22^ We employed Gαi2 in these experiments because this Gi subtype is expressed the most robustly and consistently across the cardiovascular system.^23–26^ In HEK293 suspension cells transiently transfected with human AT1R and the G protein biosensors, AngII activated Gi with approximately 2-fold lower maximal efficacy (E_max_) and 5-fold lower potency (EC_50_) than Gq **(Figure 1A, Table S1).**

**Figure 1.**
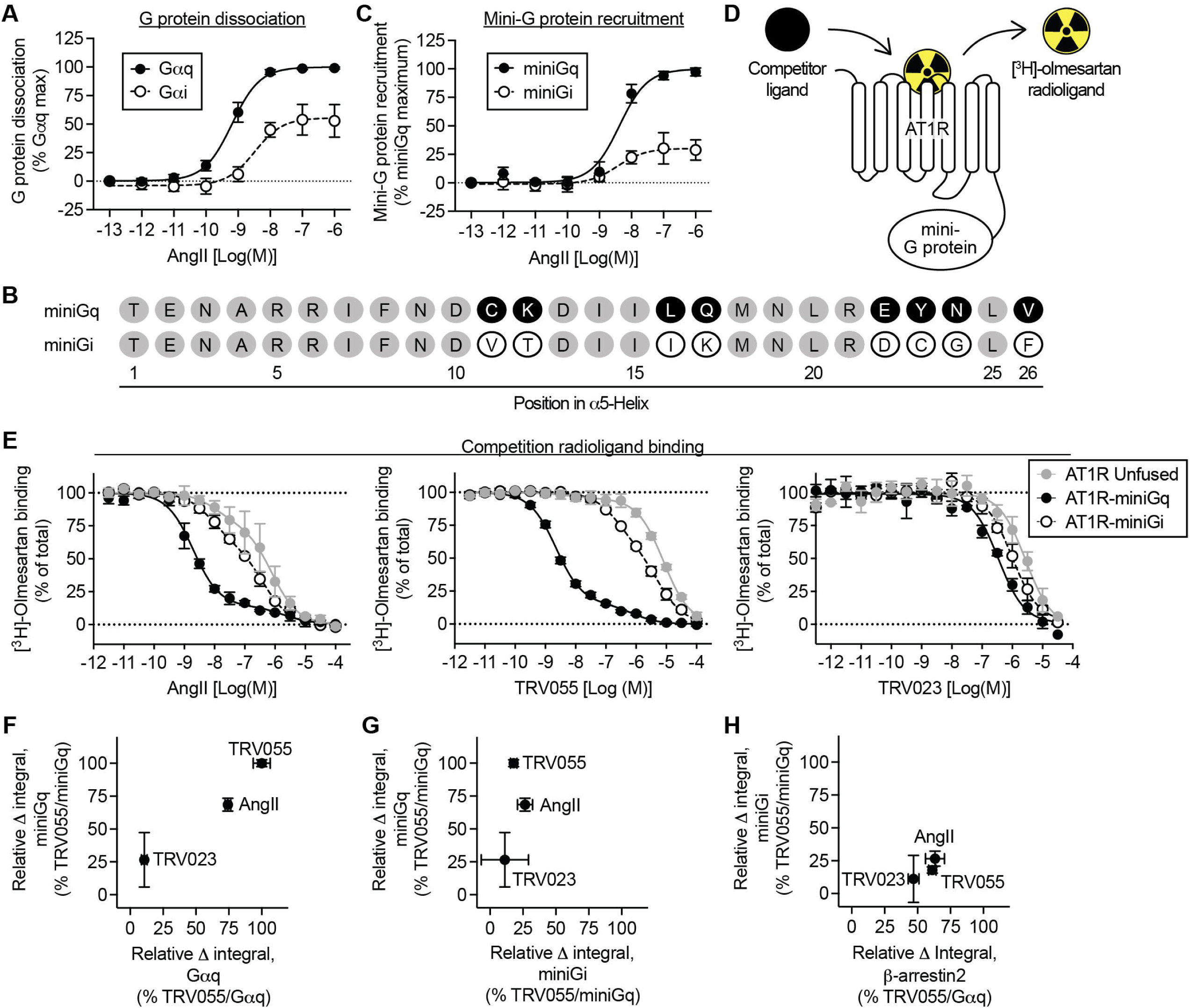
AT1R differentially activates and couples to Gq and Gi constructs. **A)** AngII-induced activation of heterotrimeric Gq and Gi in transiently transfected HEK suspension cells, monitored by decreased BRET between Gα-RLuc8 and Gγ-GFP2. Data represent the mean ± S.D. of 3 independent experiments each normalized to the Gαq E_max_ in that experiment (100%). Curve fit parameters are provided in Table S1. **B)** Alignment of the C-terminal α5 helices of miniGq and miniGi. **C)** AngII-induced recruitment of miniGq- and miniGi-mVenus to AT1R-RLuc8 in transiently transfected HEK suspension cells. Data represent the mean ± S.D. of 3 independent experiments each normalized to the miniGq E_max_ in that experiment (100%). Curve fit parameters are provided in Table S2. **D)** Schematic of AT1R-miniG fusion proteins used for radioligand binding. **E)** Competition of AngII, TRV055, and TRV023 with [^3^H]-olmesartan in membranes prepared from suspension HEK cells overexpressing the indicated AT1R-miniG fusion. Data represent the mean ± S.D. of 3 independent experiments each normalized to the top and bottom of the competition curve. **F-H)** Comparison of allosteric shifts (relative Δ integral) induced by the indicated AT1R-transducer fusions and ligands. Values for miniGq and miniGi are from panel E and normalized to TRV055/miniGq (100%); values for Gαq and β-arrestin2 are from reference 28 and normalized to that of TRV055/Gαq (100%). Data represent the mean ± S.D. of 3 independent experiments.

We next sought to identify the regions of Gα that contribute to these differences in activation by AT1R. While Gαq and Gαi2 share only 41% homology at the amino acid level, the 26-amino acid C-terminal α5 helix of Gα subunits is known to be a critical determinant of GPCR/G protein coupling specificity. Mini-G proteins, the engineered GTPase domains of Gα subunits,^16^ provide a system for testing the sufficiency of this region. MiniGq and miniGi consist of several residues from the Gαq and Gαi1/2 C-terminus, respectively, grafted onto the mini-G protein scaffold derived from Gαs; thus, they differ by only 8 residues overall **(Figure 1B, Figure S1)**. (To facilitate comparisons among constructs, residues are numbered according to their position in the 26-amino acid α5 helix rather than in the full-length proteins.) These two mini-G proteins recapitulate the receptor specificity of Gq and Gi.^16^ We compared the ability of AT1R-RLuc to recruit mVenus-tagged mini-G proteins following AngII stimulation using a BRET-based proximity assay.^20^ AngII stimulates miniGi recruitment to AT1R with approximately 3-fold lower efficacy than miniGq **(Figure 1C, Table S2)**, pointing to the key role of the Gα α5 helix in this process.

Next, we asked whether these differences in subtype-specific G protein activation and recruitment correlate with differences in allosteric coupling between AT1R and G proteins. Agonist binding to a GPCR results in conformational changes that increase the receptor’s affinity for G proteins.^27^ Reciprocally, G proteins increase the receptor’s affinity for agonists. These allosteric effects can be observed by radioligand binding, where addition of a G protein causes a leftward shift in the displacement curve of an unlabeled agonist competing with a radiolabeled antagonist.

To ensure quantitative coupling of AT1R to the G protein of interest, we fused miniGq and miniGi to the native C-terminus of AT1R via a flexible (GGGSG)_2_ linker **(Figure 1D)**, a strategy that was previously applied to full-length Gα subunits.^28,29^ To validate the construct design, we assessed the ligand dependence of allosteric shifts in [^3^H]-olmesartan competition curves at AT1R-miniGq in HEK suspension cell membranes. Compared to unfused AT1R, AT1R-miniGq showed a substantial leftward shift in the AngII competition curve **(Figure 1E)**. MiniGq led to an even more robust shift for TRV055, an AngII analog with enhanced coupling to Gq,^28,30^ but it had only a modest effect on the affinity of TRV023, a β-arrestin-biased AngII analog which does not appreciably couple to Gq.^13,28,30^ Importantly, saturation binding showed that the fusion does not alter AT1R affinity for [^3^H]-olmesartan, an antagonist **(Figure S2, Table S3)**. We quantified the allosteric shifts for AngII and its analogs by determining the area between the unfused AT1R and AT1R-miniGq curves (relative Δ integral). These values correlate well with those previously reported for these ligands at AT1R fused to full-length Gαq **(Figure 1F)**.^28^

We then compared the allosteric effects of miniGq and miniGi fusions. For all three ligands, the miniGi fusion produced a smaller leftward shift in the competition curve than miniGq **(Figure 1E)**. In contrast to miniGq, the allosteric shifts for miniGi did not show significant variation among ligands **(Figure 1G)**. This pattern of allosteric coupling for miniGi is similar to that previously observed for β-arrestin2 **(Figure 1H)**.^28^ Together these results suggest that differences in AT1R’s efficacy in Gq and Gi activation may originate in the strength of agonist-dependent allosteric coupling.

### Four α5 helix residues drive the differential coupling of Gαq and Gαi2

Since miniGq and miniGi differ only by 8 amino acids in the α5 helix, we investigated which residues contribute to the differential allosteric coupling of these two constructs. Using our AT1R-miniGi fusion, we systematically replaced each miniGi-specific residue with the corresponding miniGq residue and performed competition radioligand binding with TRV055. We employed TRV055 in these experiments since miniGq has the largest effect on its affinity, thus maximizing the ability of the assay to detect small changes. Four substitutions in the miniGi background—α5 I16L, K17Q, C23Y, and F26V—substantially potentiated the allosteric effects of miniGi by 1.8- to 2.6-fold **(Figure 2A**, **2B)**. Combining these four substitutions (miniGi α5 I16L K17Q C23Y F26V) was sufficient to generate a construct with coupling approaching that of miniGq **(Figure 2A,B, Table S4)**.

**Figure 2.**
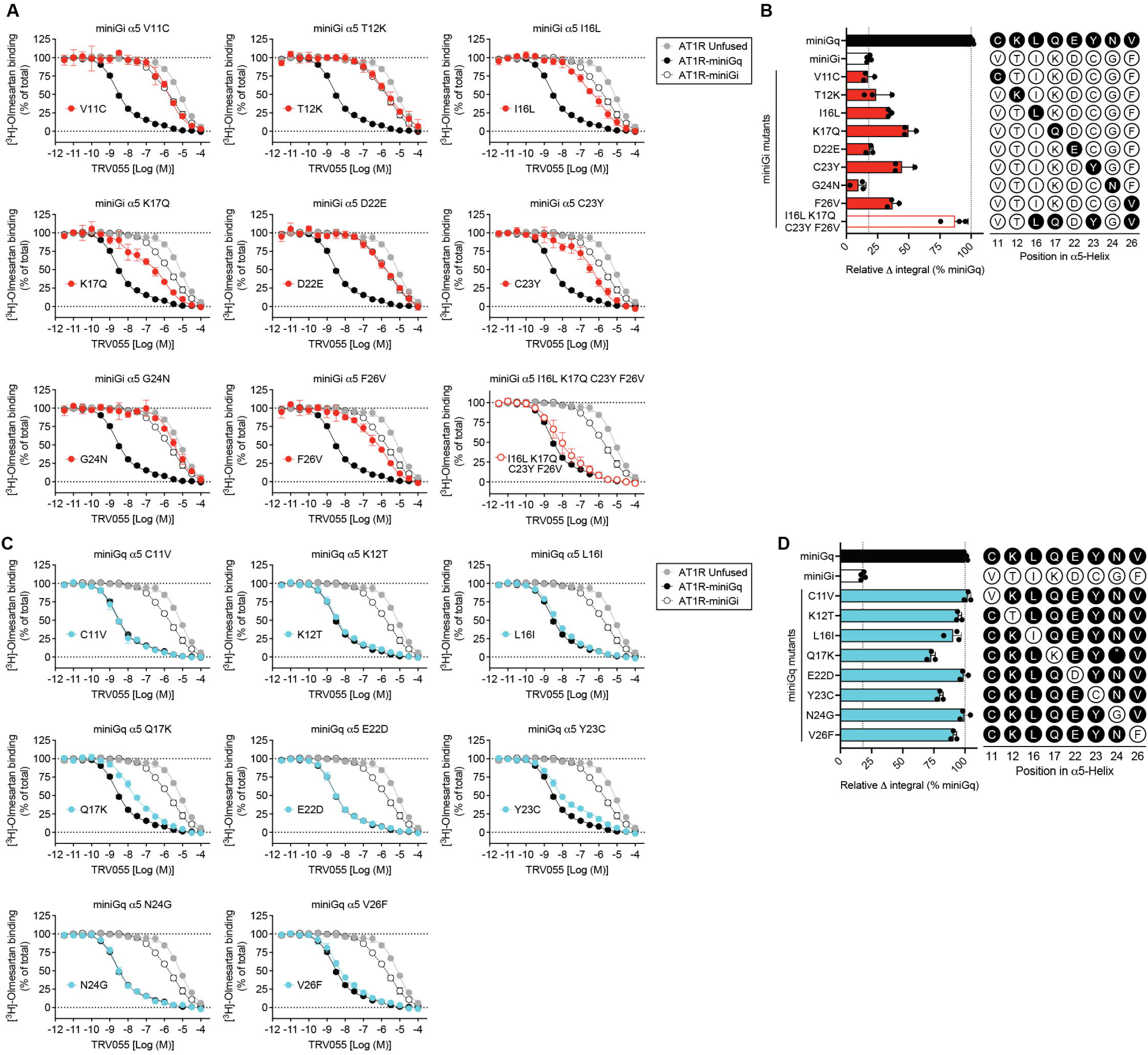
Contributions of Gq- and Gi-specific α5 helix residues to allosteric coupling to AT1R. **A,C)** Competition binding between TRV055 and [^3^H]-olmesartan in membranes prepared from HEK suspension cells overexpressing the indicated AT1R-miniG constructs. Control curves for AT1R unfused, AT1R-miniGq, and AT1R-miniGi are reproduced across panels for comparison. Data represent the mean ± S.D. of 3 independent experiments each normalized to the top and bottom of the competition curve. **B, D)** Allosteric shifts (relative Δ integral values, normalized to 100% of miniGq) for the data shown in panels A and C. Data represent the mean ± S.D. of 3 independent experiments. Means and 95% CIs are reported in Table S4.

The reciprocal swap was also performed in the miniGq background, replacing individual residues with the corresponding miniGi residue. The miniGq α5 Q17K and Y23C substitutions displayed the greatest loss of function, with allosteric effects (relative Δ integral values) 73% (95% CI: 65–81%) and 80% (95% CI: 74–86%) of miniGq, respectively. L16I and V26F had smaller effects, reducing allosteric coupling by ∼10-13% (**Figure 2C,D, Table S4**).

We next asked whether the Gαi2- and Gαq-specific residues in the α5 helix that drive differential allosteric coupling *in vitro* also account for the differential activation of these two G proteins in a cellular context. First, we tested miniGi α5 I16L K17Q C23Y F26V in the previously described mini-G protein recruitment assay. When stimulated with a dose response of TRV055, the quadruple mutant restored efficacy to the level of miniGq **(Figure 3A,B, Table S5)**. We then added these four substitutions to full-length Gαi2 and assessed TRV055-stimulated G protein dissociation. Here again, the quadruple mutant fully rescued the impaired potency and efficacy of Gαi2 **(Figure 3C,D, Table S6)**. These data suggest that these four residues in the α5 helix are sufficient to account for subtype-specific differences in AT1R activation of Gq and Gi in a cellular context.

**Figure 3.**
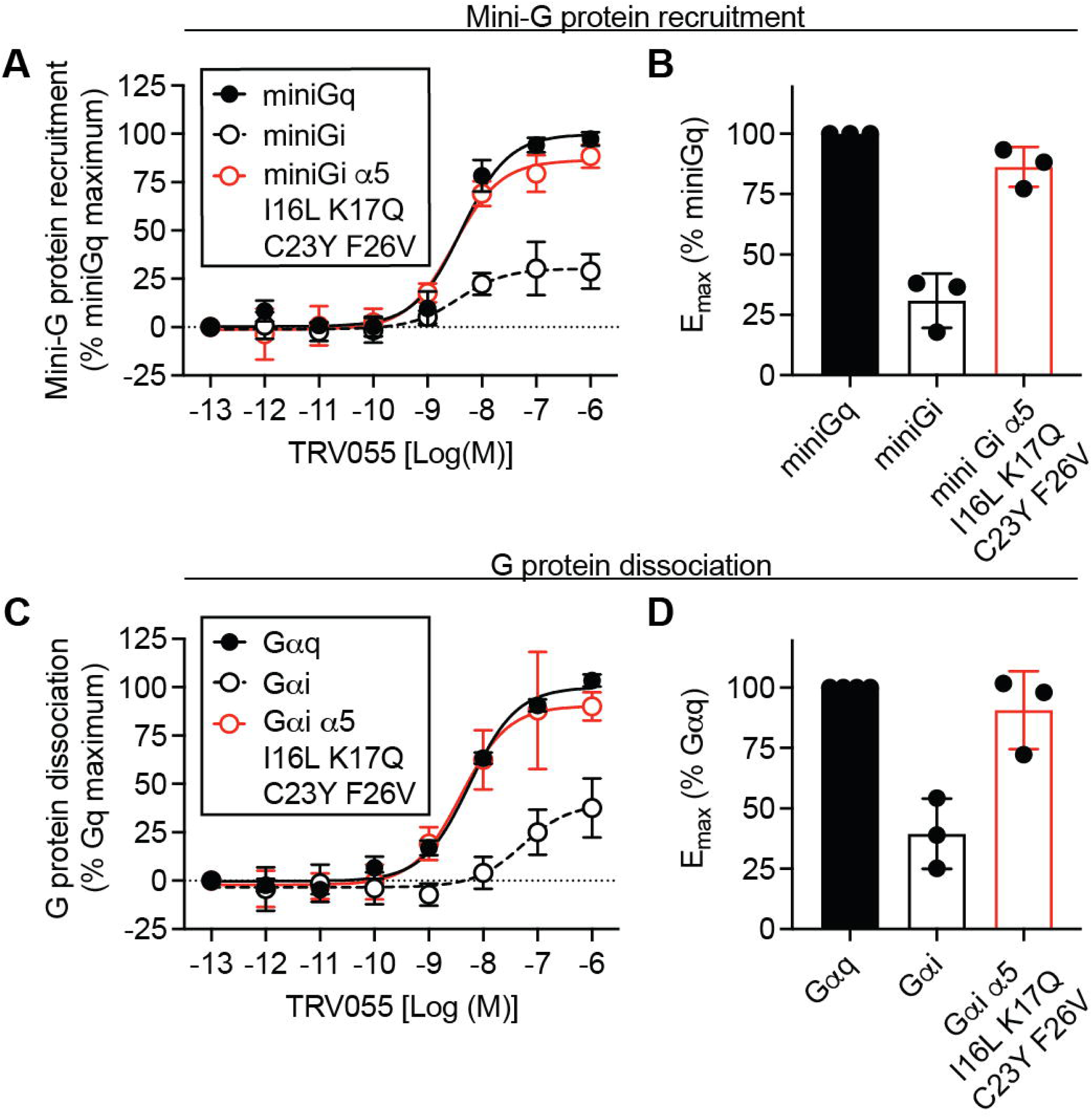
Rescue of reduced Gi activation in cellular assays by four Gq-specific substitutions in the α5 helix. **A)** TRV055-induced recruitment of miniGi α5 mutants to AT1R in transiently transfected HEK suspension cells, monitored by increased BRET between AT1R-RLuc8 and miniG-mVenus. Data represent the mean ± S.D. of 3 independent experiments each normalized to the miniGq E_max_ in that experiment (100%). **B)** E_max_ values for recruitment of mini-G protein constructs, normalized to miniGq (100%). Data represent the mean ± S.D. of 3 independent experiments. Curve fit parameters are provided in Table S5. **C)** TRV055-induced dissociation of heterotrimeric G proteins in transiently transfected HEK suspension cells, monitored by the decreased BRET between Gα-RLuc8 and Gγ-GFP2. Data represent the mean ± S.D. of 3 independent experiments each normalized to the Gαq E_max_ (100%) in that experiment. **D)** E_max_ values for G protein dissociation, normalized to Gαq (100%). Data represent the mean ± S.D. of 3 independent experiments. Curve fit parameters are provided in Table S6.

### MiniGq has higher affinity than miniGi for AT1R

We then sought to better understand the mechanism of differential coupling of Gq and Gi to AT1R. The difference in allosteric coupling strength between miniGq and miniGi observed by radioligand binding could result from lower-affinity interactions between AT1R and miniGi. Alternatively, miniGi could interact with the intracellular regions of AT1R in a manner that does not effectively promote allosteric changes in the ligand-binding site. To distinguish between these possibilities, we used Systematic Protein Affinity Strength Modulation (SPASM) biosensors.^18,19,31–33^ These constructs consist of AT1R tethered to a mini-G protein^18^ via an ER/K α-helix linker flanked by two halves of a NanoBit split luciferase^19^ **(Figure 4A)**. The rigid ER/K linker modulates the effective concentration of miniG. A sufficiently strong interaction between AT1R and miniG brings the NanoBit fragments into close proximity, reconstituting a functional luciferase enzyme.

**Figure 4.**
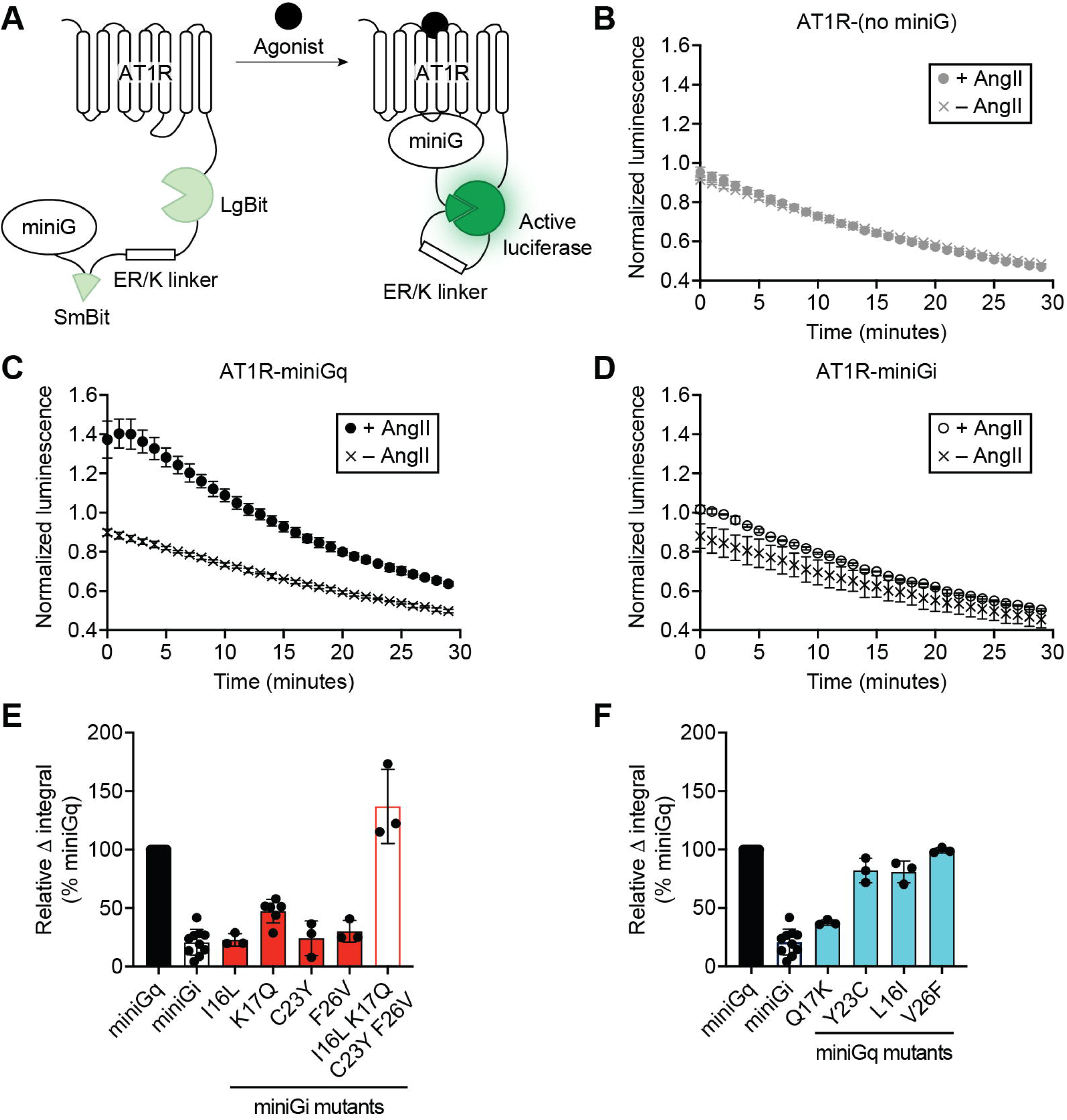
Differential interaction of miniGq and miniGi with AT1R detected with SPASM biosensors. **A)** AT1R-mini-G protein SPASM biosensor design. Agonist-dependent interactions between AT1R and mini-G proteins promote reconstitution of a functional luciferase from the LgBit and SmBit fragments. **B-D)** Luminescence time course of the indicated SPASM biosensors in the absence and presence of AngII. Data are normalized to baseline luminescence readings collected before the addition of ligand. The mean ± S.D. of 3 technical replicates from one representative experiment is shown. **E,F)** Quantification of the AngII-dependent change in the area under the time course curves (relative Δ integral) for AT1R-miniG SPASM biosensors with the indicated α5 helix mutations. Data represent the mean ± S.D. from 3-10 independent experiments and are normalized to miniGq (100%). Means and 95% CIs are reported in Table S7.

To compare the interaction of miniGq and miniGi with AT1R, we monitored the AngII-dependent increase in luciferase activity for control (no miniG), AT1R-miniGq, and AT1R-miniGi SPASM sensors in giant plasma membrane vesicles (GPMVs) prepared from HEK suspension cells overexpressing each construct. GPMV vesiculation enriches receptors that reach the plasma membrane and are fully accessible to membrane-impermeable ligands.^34^ Addition of AngII results in a robust increase in luciferase activity for the miniGq sensor but very little response for the miniGi or control constructs **(Figure 4B-D)**. The strength of the response was quantified by determining the area between the no ligand and +AngII time course curves for each construct and normalizing this value to that of miniGq (relative Δ integral) **(Figure 4E)**.

We performed reciprocal mutagenesis in the SPASM sensors at the four individual positions that contribute to allosteric coupling. Only miniGi α5 K17Q (relative Δ integral 49% of miniGq, 95% CI: 38–59%) showed increased interaction in this assay compared to miniGi (20%, 95% CI: 13–28%) **(Figure 4E, Table S7)**. This is consistent with mini Gi α5 K17Q having the largest effect in radioligand binding **(Figure 2A,B)**. However, as in other assays, combining all four substitutions (miniGi α5 I16L K17Q C23Y F26V) rescued interaction to the level of miniGq **(Figure 4E)**. This demonstrates that the other three residues collectively contribute to affinity, even though their individual contributions cannot be detected in this assay. Likewise, miniGq α5 Q17K (37%, 95% CI: 32–43%) showed greatly reduced interaction with AT1R compared to miniGq, while trends toward reduced interaction for miniGq α5 L16I and Y23C did not reach significance **(Figure 4F, Table S7)**. While the SPASM sensors appear to be less sensitive to the detection of small changes in coupling than radioligand binding, these data confirm that the impact of α5 helix substitutions on allosteric coupling is principally due to effects on the extent rather than on the mode of AT1R-miniG interaction.

### Interactions with AT1R ICL3 drive the effect of α5 residue 17 on allosteric coupling

Since α5 residue 17 has the greatest impact on coupling in both radioligand binding and SPASM assays, we sought to identify the AT1R residues that mediate its effects. We examined the cryo-EM structure of AT1R-Gq,^35^ which did not show any interactions between AT1R and Gq α5 Q17 that would rationalize the detrimental effects of the miniGq α5 Q17K substitution (**Figure 5A,B**). Based on prior simulations of AT1R^36^ and the proximity of Gq α5 Q17 to AT1R intracellular loop 3 (ICL3), a region that is often dynamic in GPCRs, we hypothesized that Q17K could create electrostatic repulsion with charged residues in ICL3, such as AT1R K232. Consistent with this hypothesis, an AT1R K232Q substitution in the context of miniGq α5 Q17K restored coupling to levels of the wild-type fusion **(Figure 5C, Table S8)**. In addition, an α5 Q17E single mutation, which introduces a negative rather than a positive charge at this position, did not reduce miniGq coupling. Together, these data suggest that the adverse impact of lysine at α5 position 17 is due to unfavorable interactions with AT1R K232.

**Figure 5.**
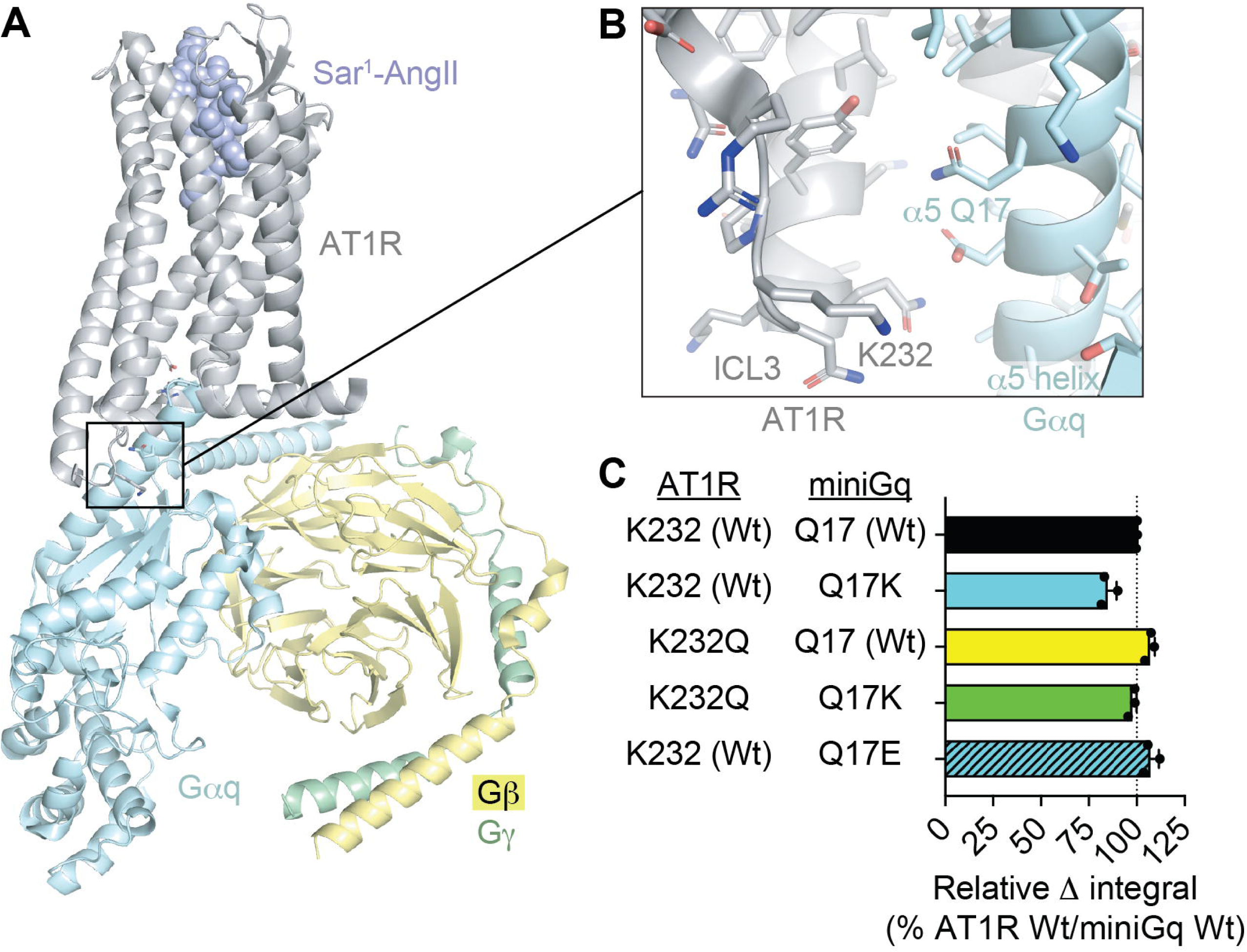
Interactions of Gαq α5 Q17. **A,B)** Structural model of AT1R bound to the agonist Sar^1^-AngII and heterotrimeric Gq determined by cryo-EM (PDB ID: 7F6G), highlighting Gαq α5 Q17 (panel B). **C)** Allosteric shifts (relative Δ integral values) in competition binding experiments between TRV055 and [^3^H]-olmesartan for AT1R-miniGq fusions with the indicated mutations. Data represent the mean ± S.D. of 3 independent experiments and are normalized to AT1R K232 (Wt)/miniGq Q17 (Wt) (100%). Means and 95% CIs are reported in Table S8.

## DISCUSSION

Many Gq-coupled GPCRs, including the AT1R, utilize Gi as a secondary G protein partner.^1,2^ In this study, we homed in on the specific structural elements and steps of G protein activation that drive differences in the coupling of these two G protein subtypes to AT1R. The use of miniG proteins for much of our work comes with limitations, since these constructs contain stabilizing mutations and deletions of the N-terminal α-helix and α-helical domain.^16^ However, since miniGq and miniGi diverge from miniGs only in the α5 helix, positions not conserved among Gα subunits are derived from Gαs. Since Gαs does not couple to AT1R^2,37^, the mini-G protein backbone provides a “neutral” scaffold. Therefore, miniGq and miniGi are valuable tools that demonstrate the sufficiency of a limited number of subtype-specific residues to recapitulate differences in the coupling strength of Gαq and Gαi2 to AT1R **(Figure 1)**. We showed that replacing four miniGi residues (α5 16, 17, 23, and 26) with Gαq-specific residues fully restores the strong allosteric coupling found in miniGq **(Figure 2)**. These four substitutions in Gαi2 are also sufficient to rescue the reduced activity of Gi in both miniG protein recruitment and heterotrimeric G protein activation **(Figure 3)**, attesting to their importance in a cellular context. Our findings are consistent with previous work showing that replacing residues α5 17-26 of Gαi1 with those from Gαq yields a chimera that can be recruited to AT1R almost as robustly as Gαq,^2^ and they enhance our understanding of the underlying mechanisms.

There are now multiple examples of Family A (rhodopsin-like) GPCRs whose structures have been determined in complex with both Gq and Gi family members, or with the corresponding mini-G proteins. Some of these matched pairs show substantial differences in the receptor’s conformation and in the positioning of the Gα α5 helix (e.g., references 38-40). Other complexes have very similar overall architectures at the GPCR-Gα interface and principally vary in specific contacts (e.g., references 41-43). Our data show that replacement of four different Gαi2-specific residues with those found in Gαq can independently increase miniGi coupling to AT1R **(Figure 2A,B)**. This suggests that miniGq and miniGi engage the AT1R 7TM core in a similar manner, such that major conformational changes are not required to restore the AT1R contacts responsible for miniGq’s strong coupling. Notably, only two of the four residues we identified are within the “wavy hook,” the 4-5 residues at the distal end of the Gα α5 helix that form an α-helical turn upon receptor engagement. This region has received the most attention as a driver of Gα specificity^44^ and GPCR engagement of primary versus secondary G proteins.^45^

Structural data have highlighted the critical role of GPCR ICLs in G protein activation and recognition. We present evidence that the critical contribution of α5 residue 17, which has the largest effect on allosteric coupling and interaction with AT1R **(Figures 2**, **4E,F)**, is mediated by interactions with AT1R ICL3 **(Figure 5)**. While Gαq α5 Q17 interacts with the ICL3 of several other GPCRs,^42,46^ direct contacts with ICL3 were not observed in the AT1R-Gq structure.^35^ Therefore, the impact of α5 Q17 on miniGq’s strong allosteric coupling and its interaction with ICL3 would have been difficult to predict from the structure alone. This highlights the importance of using functional assays and biophysical and computational approaches that capture protein dynamics,^36^ rather than structures alone, to understand patterns of G protein coupling to GPCRs.

Interestingly, AngII, Gq-biased, and β-arrestin-biased ligands all exhibit comparable allosteric coupling to miniGi, just as they have comparable allosteric coupling to β-arrestin **(Figure 1E,G)**.^28^ This observation accords with previous reports that many β-arrestin-biased AT1R ligands robustly activate Gi.^47,48^ These data suggest that despite being structurally unrelated, Gi and β-arrestin are both able to engage the full range of AT1R conformations that these ligands stabilize.^49^

In conclusion, we found that a finite number of residues dispersed through the distal half of the Gα α5 helix account for differences in the strength of AT1R’s coupling to Gq and Gi at each phase of G protein activation, from ligand-induced allosteric interaction to heterotrimer dissociation. Across the broader GPCR family, there appears to be substantial diversity in what structural elements are required for GPCRs to maximally activate G proteins. While some receptors require only the most distal residues of the Gα α5 helix,^2,14^ others are affected by elements outside of this region.^2,50^ A better understanding of the molecular mechanisms that determine the strength of coupling should ultimately guide the development of pharmacological approaches to selectively modulate the activation of each G protein subtype.

## DATA AVAILABILITY STATEMENT

The authors declare that all the data supporting the findings of this study are available within the paper and its Supplemental Material.

## CRediT AUTHORSHIP CONTRIBUTION STATEMENT

Akua A. Acheampong: Conceptualization, Formal analysis, Funding acquisition, Investigation, Methodology, Resources, Visualization, Writing – original draft, Writing – review and editing; **Matthew K. Collins:** Methodology; **Amanda R. Keller:** Methodology; **Andrew P. Feld:** Resources; **Brandi M. Small:** Resources; **Michael Ritt:** Methodology, Resources; **Carl-Mikael Suomivuori:** Funding acquisition, Methodology; **Sivaraj Sivaramakrishnan:** Funding acquisition, Methodology, Resources; **Laura M. Wingler:** Conceptualization, Formal analysis, Funding acquisition, Investigation, Methodology, Resources, Supervision, Visualization, Writing – original draft, Writing – review and editing

## Non-standard abbreviations

AngII: angiotensin II
AT1R: angiotensin II type 1 receptor
AUC: area under the curve
BRET: bioluminescence resonance energy transfer
CI: confidence interval
E_max_: maximal efficacy
GFP2: mutant green fluorescence protein
GPCR: G protein-coupled receptor
GPMV: giant plasma membrane vesicle
ICL3: intracellular loop 3
LgBit: Large Bit Nanoluciferase fragment
PBS: phosphate-buffered saline
Rluc8: mutant *Renilla* luciferase
SmBit: Small Bit Nanoluciferase fragment
SPASM: Systematic Protein Affinity Strength Modulation
Wt: wild-type

