## Supplementary materials for "Molecular Determinants of G Protein Subtype-Specific Coupling to the Angiotensin II Type 1 Receptor"

**Contents:**

Tables S1 to S8
Figures S1 to S2

**Molecular Determinants of G Protein Subtype-Specific Coupling to the Angiotensin II Type 1 Receptor**

Akua A. Acheampong, Matthew K. Collins, Amanda R. Keller, Andrew P. Feld, Brandi M. Small, Michael Ritt, Carl-Mikael Suomivuori, Sivaraj Sivaramakrishnan, Laura M. Wingler

**Table S1**

| Construct | $LogEC_{50}$ [95% CI] | Relative $E_{max}$ (% of Gαq) | n |
|---|---|---|---|
| Gαq | -9.2 [-9.6, -8.8] | 100 | 3 |
| Gαi2 | -8.5 [-9.0, -8.0] | 56 [21, 90] | 3 |

Data represent the mean $logEC_{50}$ and $E_{max}$ values from individual experiments and corresponding 95% confidence intervals for AngII-induced activation of heterotrimeric Gq and Gi by AT1R.

**Table S2**

| Construct | $LogEC_{50}$ [95% CI] | Relative $E_{max}$ (% of miniGq) | n |
|---|---|---|---|
| miniGq | -8.4 [-8.8, -8.0] | 100 | 3 |
| miniGi | -8.5 [-10.0, -7.0] | 31 [3, 59] | 3 |

Data represent the mean $logEC_{50}$ and $E_{max}$ values from individual experiments and corresponding 95% confidence intervals for AngII-induced recruitment of mini-G proteins by AT1R.

**Table S3**

| Construct | Kd (nM) [95% CI] | n |
|---|---|---|
| AT1R (Unfused) | 0.58 [0.51, 0.66] | 3 |
| AT1R-miniGq | 0.66 [0.41, 0.91] | 3 |
| AT1R-miniGi | 0.64 [0.47, 0.82] | 3 |

Data represent the mean Kd values and corresponding 95% confidence intervals for [$^3$H]-olmesartan binding to suspension HEK cell membranes overexpressing the indicated constructs determined by saturation binding.

**Molecular Determinants of G Protein Subtype-Specific Coupling to the Angiotensin II Type 1 Receptor**

Akua A. Acheampong, Matthew K. Collins, Amanda R. Keller, Andrew P. Feld, Brandi M. Small, Michael Ritt, Carl-Mikael Suomivuori, Sivaraj Sivaramakrishnan, Laura M. Wingler

**Table S4**

| Construct | | Relative Δ integral (% miniGq) [95% CI] | n |
|---|---|---|---|
| miniGq | | 100<br>[97, 103] | 4 |
| miniGi | | 18<br>[15, 21] | 4 |
| miniGi background | V11C | 17<br>[6, 29] | 3 |
| | T12K | 24<br>[-5, 52] | 3 |
| | I16L | 35<br>[30, 39] | 3 |
| | K17Q | 50<br>[37, 63] | 3 |
| | D22E | 19<br>[12, 26] | 3 |
| | C23Y | 45<br>[22, 67] | 3 |
| | G24N | 10<br>[-5, 25] | 3 |
| | F26V | 37<br>[25, 49] | 3 |
| | I16L K17Q C23Y F26V | 87<br>[61, 113] | 3 |
| miniGq background | C11V | 102<br>[96, 109] | 3 |
| | K12T | 95<br>[89, 101] | 3 |
| | L16I | 90<br>[74, 107] | 3 |
| | Q17K | 73<br>[65, 81] | 3 |
| | E22D | 99<br>[90, 108] | 3 |
| | Y23C | 80<br>[74, 86] | 3 |
| | N24G | 99<br>[89, 110] | 3 |
| | V26F | 90<br>[85, 97] | 3 |

Relative Δ integral values for allosteric shifts induced by the indicated AT1R-fusion proteins in competition binding experiments between TRV055 and [$^3$H]-olmesartan. Residues are numbered according to their position in the α5 helix.

**Molecular Determinants of G Protein Subtype-Specific Coupling to the Angiotensin II Type 1 Receptor**

Akua A. Acheampong, Matthew K. Collins, Amanda R. Keller, Andrew P. Feld, Brandi M. Small, Michael Ritt, Carl-Mikael Suomivuori, Sivaraj Sivaramakrishnan, Laura M. Wingler

**Table S5**

| Construct | $LogEC_{50}$ [95% CI] | Relative $E_{max}$ (% of miniGq) | n |
|---|---|---|---|
| miniGq | -8.4 [-8.8, -8.0] | 100 | 3 |
| miniGi | -8.5 [-10 -7.0] | 31 [3, 59] | 3 |
| miniGi α5 I16L K17Q C23Y F26V | -8.5 [-8.7, -8.3] | 86 [66, 107] | 3 |

Data represent the mean $logEC_{50}$ and $E_{max}$ values from individual experiments and corresponding 95% confidence intervals for TRV055-induced recruitment of mini-G proteins by AT1R. Residues are numbered according to their position in the α5 helix.

**Table S6**

| Construct | $LogEC_{50}$ [95% CI] | Relative $E_{max}$ (% of Gq) | n |
|---|---|---|---|
| Gαq | -8.2 [-8.4, -8.1] | 100 | 3 |
| Gαi2 | -7.2 [-8.0, -6.5] | 39 [3, 76] | 3 |
| Gαi2 α5 I16L K17Q C23Y F26V | -8.4 [-9.1, -7.8] | 91 [51, 131] | 3 |

Data represent the mean $logEC_{50}$ and $E_{max}$ values from individual experiments and corresponding 95% confidence intervals for TRV055-induced activation of heterotrimeric G proteins with the indicated Gα subunits. Residues are numbered according to their position in the α5 helix.

**Molecular Determinants of G Protein Subtype-Specific Coupling to the Angiotensin II Type 1 Receptor**

Akua A. Acheampong, Matthew K. Collins, Amanda R. Keller, Andrew P. Feld, Brandi M. Small, Michael Ritt, Carl-Mikael Suomivuori, Sivaraj Sivaramakrishnan, Laura M. Wingler

**Table S7**

<table>
<tr><th colspan="2">SPASM Construct</th><th>Relative $\Delta$ integral<br>(% miniGq)<br>[95% CI]</th><th>n</th></tr>
<tr><td colspan="2">miniGq</td><td>100</td><td>10</td></tr>
<tr><td colspan="2">miniGi</td><td>20<br>[13, 28]</td><td>10</td></tr>
<tr><td rowspan="5">miniGi background</td><td>I16L</td><td>23<br>[10, 36]</td><td>3</td></tr>
<tr><td>K17Q</td><td>49<br>[38, 59]</td><td>3</td></tr>
<tr><td>C23Y</td><td>24<br>[-13, 61]</td><td>3</td></tr>
<tr><td>F26V</td><td>30<br>[7, 53]</td><td>3</td></tr>
<tr><td>I16L K17Q<br>C23Y F26V</td><td>137<br>[58, 216]</td><td>3</td></tr>
<tr><td rowspan="4">miniGq background</td><td>L16I</td><td>81<br>[58, 104]</td><td>3</td></tr>
<tr><td>Q17K</td><td>37<br>[32, 43]</td><td>3</td></tr>
<tr><td>Y23C</td><td>82<br>[56, 108]</td><td>3</td></tr>
<tr><td>V26F</td><td>99<br>[94, 105]</td><td>3</td></tr>
</table>

Relative $\Delta$ integral values and corresponding 95% confidence intervals for the AngII-induced change in normalized luminescence for the indicated AT1R-miniG SPASM biosensors. Residues are numbered according to their position in the $\alpha$5 helix.

**Molecular Determinants of G Protein Subtype-Specific Coupling to the Angiotensin II Type 1 Receptor**

Akua A. Acheampong, Matthew K. Collins, Amanda R. Keller, Andrew P. Feld, Brandi M. Small, Michael Ritt, Carl-Mikael Suomivuori, Sivaraj Sivaramakrishnan, Laura M. Wingler

**Table S8**

| **AT1R** | **miniGq** | **Relative Δ integral (% miniGq) [95% CI]** | **n** |
|---|---|---|---|
| K232 (Wt) | Q17 (Wt) | 100<br>[99, 101] | 3 |
| K232 (Wt) | Q17K | 85<br>[73, 96] | 3 |
| K232Q | Q17 (Wt) | 107<br>[101, 113] | 3 |
| K232Q | Q17K | 98<br>[93, 103] | 3 |
| K232 (Wt) | Q17E | 107<br>[96, 118] | 3 |

Relative Δ integral values for allosteric shifts induced by the indicated AT1R-fusion proteins in competition binding experiments between TRV055 and [$^3$H]-olmesartan. Residue numbering for miniGq reflects the position in the $\alpha$5 helix.

**Molecular Determinants of G Protein Subtype-Specific Coupling to the Angiotensin II Type 1 Receptor**

Akua A. Acheampong, Matthew K. Collins, Amanda R. Keller, Andrew P. Feld, Brandi M. Small, Michael Ritt, Carl-Mikael Suomivuori, Sivaraj Sivaramakrishnan, Laura M. Wingler

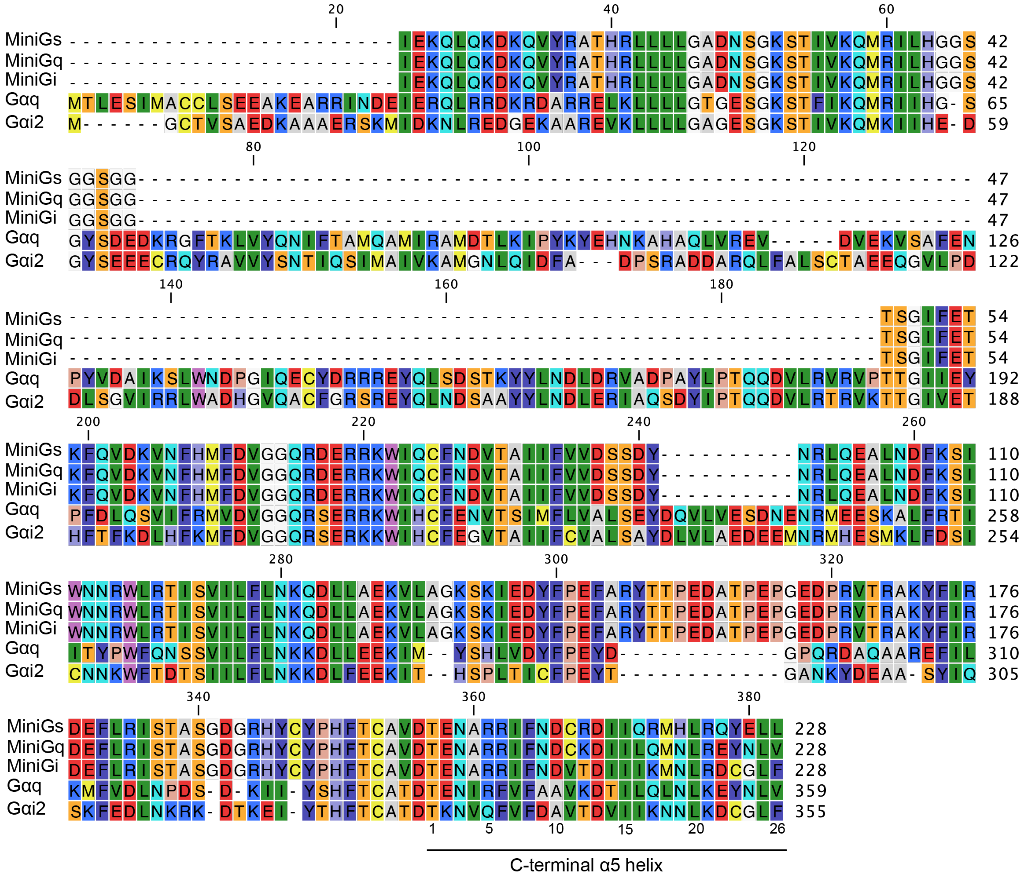


**Figure S1. Alignment of miniGs, miniGq (miniGs/q70), miniGi (miniGs/i43), Gαq, and Gαi2 protein sequences**.

**Molecular Determinants of G Protein Subtype-Specific Coupling to the Angiotensin II Type 1 Receptor**

Akua A. Acheampong, Matthew K. Collins, Amanda R. Keller, Andrew P. Feld, Brandi M. Small, Michael Ritt, Carl-Mikael Suomivuori, Sivaraj Sivaramakrishnan, Laura M. Wingler

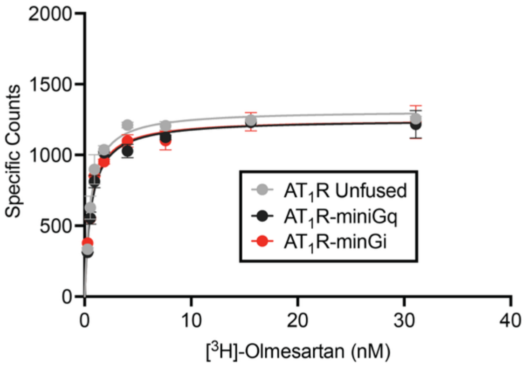


**Figure S2. Saturation binding of [$^{3}$H]-Olmesartan to HEK suspension cell membranes overexpressing the indicated AT1R constructs.** The mean ± S.D. of two technical replicates from a single representative experiment is shown. The $K_d$ values (mean and 95% confidence intervals) determined from three independent experiments are reported in Table S3.
